# FLASH radiotherapy spares lymphocytes in tumor-draining lymph nodes and increases infiltration of immune cells in tumors

**DOI:** 10.1101/2025.04.07.647544

**Authors:** Francisco R. Saenz, Brett Velasquez, Trey Waldrop, Edgardo Aguilar, Kathryn R Cox, Abagail Delahoussaye, Caddie Laberiano-Fernandez, Leticia Campos Clemente, Luke Connell, Nefetiti Mims, Denae Neill, Edwin R Parra, Karen Clise-Dwyer, Emil Schüler, Michael T. Spiotto

## Abstract

Radiotherapy (RT) delivered at conventional dose rates (CONV) can both stimulate antitumor immune responses and inhibit these immune responses by depleting circulating lymphocytes. Given the observed normal tissue sparing associated with ultra-high dose rate (FLASH) RT, we hypothesized that FLASH RT may protect lymphocytes while increasing the immunogenicity of cancer cells. We irradiated cancer cell lines *in vitro* with FLASH RT or CONV RT and assessed immunogenic mRNA and protein expression. Both HPV-positive cell lines MEER and TC-1 showed upregulation of *Calr, Hmgb1*, and cGAS-STING family members after FLASH RT but not after CONV RT *in vitro*. To assess changes in lymphocyte populations, we irradiated murine mEER tumors in syngeneic C57BL/6 mice with 27 Gy in 3 fractions of FLASH RT or CONV RT. In mice bearing FLASH irradiated tumors, tumor-draining lymph nodes contained greater numbers of CD8^+^ T cells (FLASH 1.7×10^4^ vs 0.8×10^4^ CONV; *P*<0.001) and CD4^+^ T cells (FLASH 2.3×10^4^ vs CONV 1.2×10^4^; *P*<0.001) after irradiation. FLASH RT was associated with increased numbers of activated CD44^+^CD62L^lo^CD8^+^ and CD4^+^ lymphocytes. In irradiated tumors, FLASH RT was associated with increased CD8^+^ tumor-infiltrating lymphocytes, increased PD1 expression on these lymphocytes and increased PDL1 expression on macrophages. Compared with CONV RT, FLASH RT spared activated T cells in tumor-draining lymph nodes and in tumors but increased checkpoint inhibitor expression in tumors. These results suggest that FLASH RT may enhance antitumor immune responses by maintaining the immunogenic effects of RT while preserving lymphocyte numbers, which may be augmented with immune checkpoint blockade.

**Significance:** Radiation-induced lymphopenia is associated with poorer survival outcomes. New treatment approaches, like FLASH radiation therapy (FLASH RT), which reduce lymphopenia and enhance the antitumor response, could potentially lead to better outcomes for cancer patients.

## Introduction

Radiotherapy (RT) stimulates immune responses against cancer by increasing cancer cell death, tumor antigen release, and antigen cross-presentation as well as causing the release of damage-associated molecular pattern proteins (DAMPs) and other immunogenic molecules that prompt maturation of dendritic cells, which in turn prime cytotoxic and helper lymphocytes in tumor draining lymph nodes (DLNs). By contrast, RT also inhibits antitumor immune responses by depleting circulating and tumor-infiltrating lymphocytes (1), by inducing dysfunction in tumor-DLNs, and by increasing immunosuppressive cells in the tumor microenvironment (2). Radiation-induced lymphopenia has been observed in several types of cancer, including cervix (3), head and neck (4), esophageal (5), and lung (6). Radiation-induced lymphopenia has also been negatively correlated with survival during RT, suggesting that the effectiveness of radiation delivered at conventional dose rates (CONV RT) may be limited by its deleterious effects on the patient’s immune system. Consequently, a pressing need exists to identify radiation approaches that can mitigate the radiation-induced lymphopenia and immune suppression associated with treatment.

Ultra-high dose rate or FLASH RT was developed to improve the therapeutic index of tumor-ablative doses by sparing normal tissues from radiation-induced damage (7). FLASH RT is delivered at dose rates in excess of 40 Gy per second (8-11), as compared with 0.01–0.5 Gy per second for CONV RT, and this ultra-high dose rate has been shown to spare normal tissues in the skin and gastrointestinal tract (12-16). The normal tissue sparing of FLASH RT has also been shown to improve cognition after brain irradiation (17,18) and decrease fibrosis after lung irradiation (9). The proposed mechanisms of FLASH RT include: (1) a rapid depletion of oxygen leading to generation of less reactive oxygen species, (2) differences in DNA damage repair between cancer cells and normal tissues, and (3) altered inflammatory responses. However, the immune-sparing benefits of FLASH RT and the immunogenic role of FLASH RT remain vastly understudied. FLASH RT may also spare antitumor immune responses compared with CONV RT by sparing antigen-presenting cells, lymphocytes, or both. FLASH RT may also cause greater cancer cell death or greater upregulation of immunogenic pathways involving DAMPs or cGAS-STING to further stimulate antitumor immune responses (19).

Here, we assessed the immune-sparing effects of FLASH RT in tumors, DLNs, and peripheral blood from a mouse model of head and neck cancer. Relative to CONV RT, FLASH RT led to increased numbers of naïve and activated CD8^+^ T lymphocytes. These immune-sparing effects were tempered by an increase in PD1^+^ CD8^+^ T cells and PDL1^+^ macrophages in the irradiated tumors. Consequently, FLASH RT was found to increase circulating effector T cells while also increasing immune checkpoint expression in the tumor microenvironment.

## Materials and Methods

### Irradiation Setup

All irradiations were conducted by using a 9-MeV electron linear accelerator capable of delivering both CONV and ultra-high dose rate (FLASH) irradiations (Mobetron, IntraOp Medical, Sunnyvale, CA) (20). The irradiation setups were calibrated by using Gafchromic film, and daily output verification and dose monitoring was performed with ion chambers and beam current transformers as previously described (21-25). CONV irradiations were conducted with a mean dose rate of 0.2 Gy/s and a dose per pulse of 0.01 Gy/pulse. The corresponding irradiation parameters for FLASH were 270 Gy/s and 3 Gy/pulse. Dose delivery and dosimetry measurements were recorded for each delivery. A full list of the measured irradiation parameters is provided in Supplementary Tables S1 and S2.

### Cell Culture

The HPV-positive murine cell lines TC1 (lung carcinoma) and mEER (orpharyngeal squamous cell carcinoma) were a kind gift from Dr. J. Sastry (MD Anderson) and maintained in complete medium as previously described (26,27). Cells were cultured up to 70% confluency and tested for mycoplasma before injection into animals. Cell suspensions were obtained with trypsin (Gibco, Thermo Fisher Scientific, 25-200-056, Waltham, MA, USA) according to the manufacturer’s recommendation, counted, and resuspended in complete medium for irradiation experiments (TC1 and mEER) or in Dulbecco’s phosphate buffered saline (DPBS, 1X) (Gibco 14190-136) for subcutaneous syngeneic injections (mEER). Cells were irradiated in suspension (1.0-1.5 × 10^6^ cells/mL).

### Animals

Eight-to twelve-week-old C57BL/6J female mice were purchased from Jackson Laboratory (Bar Harbor, ME, USA) and housed in a pathogen-free environment. Mice were injected with 2×10^6^ mEER cells subcutaneously in 0.1 mL DPBS on the right flank. Mice bearing tumors averaging 100-150 mm^3^ in volume were randomly assigned to one of three groups: CONV RT, FLASH RT, or SHAM irradiated controls. Tumors were irradiated with a hypofractionated regimen (3 fractions of 9 Gy each), and mice were killed at 5 days after irradiation for endpoint analysis (see below). Animal studies were approved and conducted in accordance with The University of Texas MD Anderson Cancer Center Institutional Animal Care and Use Committee.

### Real-Time Quantitative PCR with Reverse Transcription

Total RNA was isolated by using TRIzol (Thermo Fisher Scientific 15596028) according to the manufacturer’s protocol. Total RNA concentration was determined with a NanoDrop OneC spectrophotometer (Thermo Fisher Scientific), and 2 µg of RNA was treated with RNase-free DNase I (Thermo Fischer Scientific EN0521) in a 10-µL reaction. cDNA was synthesized by using a High-Capacity cDNA Reverse Transcription Kit (Applied Biosystems 4374966, Foster City, CA, USA) with random primers. *Calreticulin* (Mm00482936), *Hmgb1* (Mm00849805_gH), *Ifna* (Mm03030145_gH), *Mb21d1* (Mm01147496 M1), and *Tmem173* (Mm01158117_M1) were purchased from Thermo Fisher Scientific. The qRT-PCR reactions were synthesized with TaqMan Gene Expression Master Mix (Applied Biosystems 4369016) and analyzed on a QuantStudio 5 system (Applied Biosystems, software v1.5.1). *Ct* values for every gene were subtracted from their corresponding *Gapdh* (Δ*Ct*). Normalized fold expression was obtained by using the 2^-ΔΔ*Ct*^ formula.

### Immunofluorescence

Cells were rinsed twice with cold 1X-PBS and fixed in 4% paraformaldehyde (Thermo Fisher Scientific J19943-K2) in 1X-PBS for 10 minutes at room temperature. After fixation, cells were rinsed and blocked with 1% bovine serum albumin (Cell Signaling Technology 9998S, Danvers, MA, USA) in 1X-PBS for 20-30 minutes. CALR (Clone SU37-03) (Thermo Fisher Scientific PIMA532131) was diluted in blocking buffer (1:500) and incubated for 20-30 minutes at room temperature. After incubation with primary antibody, cells were rinsed and incubated with Donkey F(ab’)2 Anti-Rabbit IgG(H+L) AF-647 secondary antibody (1:1000) for 20-30 minutes at room temperature and protected from light. Cells were then rinsed, counterstained with 4’,6-diamidino-2-phenylindole (DAPI) (1:1000) for 5 minutes at room temperature, rinsed again, and 0.1 mL of PBS was added to each well before imaging. A montage image (area 0.18 cm^2^ on a 96-well plate) was acquired with a Cytation 5 Cell Imaging Multi-Mode Reader and images analyzed with Gen5, v3.10 software (BioTek Instruments, Inc., Winooski, VT, USA).

### Flow Cytometry

Peripheral blood samples were collected from mice via retro-orbital bleeding using heparinized capillaries (Fisher Scientific 22-362-566) and collected into 0.2 mL of 20-30 IU/mL heparin (Thermo Fisher Scientific A16198) at 5 days after the initial irradiation. Tumor DLNs were harvested at necropsy and transferred into a 15 mL conical tube with 5 mL PBS kept in ice, then passed through a 70-µm cell strainer (Corning 352350, Corning, NY, USA) to prepare single-cell suspensions. The cell suspensions were treated with red blood cell lysis buffer (Thermo Fisher Scientific 00-4300-54) according to the manufacturer’s recommendation. Cells were incubated with a viability dye, washed, and resuspended in staining buffer containing anti-CD16/32 (clone 93) for 10 minutes before cell-surface staining with either a lymphoid or a myeloid panel of antibodies (Supplementary Table S3). Following staining, cells were fixed with 1% paraformaldehyde (Thermo Fisher Scientific) and analyzed with an Aurora cytometer (Cytek Biosciences, Fremont, CA, USA), equipped with five lasers (355 nm, 405 nm, 488 nm, 561 nm, and 640 nm) and capable of detecting up to 64 fluorescence parameters. Data acquisition was performed using SpectroFlo software (Cytek Biosciences) and analyzed using FlowJo software v10.10.0 (FlowJo LLC, Ashland, OR, USA). Events were quantified as absolute counts per analyzed volume of resuspended blood cells. Blood volume was estimated by weighing collected samples, which were then divided into two portions for staining with different antibody panels. Following staining, half of the resuspended blood cell volume from each portion was acquired and analyzed by the cytometer. The gating strategies for the lymphoid and myeloid panels are shown in Supplementary Figure S1.

### Multiplex Immunofluorescence Staining and Image Analysis

Multiplex staining, imaging, and analysis were done by the Translational Molecular Pathology Immunoprofiling Laboratory at MD Anderson Cancer Center (28). Briefly, formalin-fixed paraffin-embedded tumors were cut into 4-μm-thick sections and then stained for the following tumor-infiltrating lymphocyte markers: CK19, CD3, CD4, CD8, PD1, PDL1, CD19, and F4-80. All the markers were stained in sequence with their respective fluorophore contained in the Opal 7 IHC kit (catalog #NEL797001KT; Akoya Biosciences, Marlborough, MA) and the individual tyramide signal amplification fluorophores Opal Polaris 480 (catalog #FP1500001KT) and Opal Polaris 780 kit (catalog #FP1501001KT, Akoya Biosciences). Slides were scanned with a Vectra Polaris 1.0.13 (Akoya Biosciences) at low magnification, 10x (1.0 µm/pixel) through the complete emission spectrum, and positive tonsil controls from the run staining were used to calibrate the spectral image scanner protocol (29). A pathologist selected all regions of interest within the tumor areas for scanning in high magnification by the Phenochart Software image viewer 1.0.12 (931 × 698 µm size at resolution 20x) to capture various elements of tissue heterogeneity. A pathologist analyzed each region of interest by using InForm 2.4.8 image analysis software (Akoya Biosciences). Marker co-localization was used to identify specific tumor and stroma compartment cell phenotypes. The densities of each cell phenotype (Supplementary Table S4) were quantified, and the final data were expressed as the number of cells/mm^2^ (29). The data were consolidated by using R studio 3.5.3 (Phenopter 0.2.2 packet; https://rdrr.io/github/akoyabio/phenoptrReports/f/, Akoya Biosciences).

## Statistical Analysis

Data were represented as means± SEM of at least three biological replicates and graphed with GraphPad v10.0.3 (Boston, MA, USA). Differences between means were assessed with Student’s *t* tests. Statistical significance was defined as *P*<0.05.

## Data availability statement

The data generated in this study are available within the article and its Supplementary Data files.

## Results

### FLASH RT Increased the Expression of Immunogenic Markers in Cancer Cells

FLASH RT, as compared with CONV RT, has been shown to reduce immunogenic markers such as cGAS-STING in normal tissues (19). Still, it remains unclear if the expression of immunogenic proteins in cancer cells differs after FLASH RT versus after CONV RT. To investigate these differences, we examined the expression of the immunogenic markers calreticulin (*Calr*) and *Hmgb1* in murine cancer cell lines TC1 and mEER treated with increasing doses (0, 6, 9, and 12 Gy) of either FLASH RT or CONV RT. Compared with CONV RT, FLASH RT significantly increased *Calr* mRNA expression in TC1 cells in a dose dependent manner (mean fold-change 0.97 vs. 4.2, *P*<0.001) and 12 Gy (1.3 vs. 4.1, *P*<0.01) and analyzed 24 h after irradiation (Fig. 1A). Similarly, FLASH RT increased *Hmgb1* mRNA expression in TC1 cells irradiated with 9 Gy (4.2 vs 1.0, *P*<0.001) and 12 Gy (4.1 vs 1.3, *P*<0.01) at 24 h after treatment compared with CONV RT (Fig. 1B). In mEER cells, FLASH RT vs CONV RT did not cause significant changes in *Calr* expression (for 9 Gy, 1.2 vs. 1.1, *P*=0.09; for 12 Gy, 1.3 vs 1.2, *P*=0.08) (Fig. 1C) or in *Hmgb1* expression (for 12 Gy, 2.5 vs. 1.2, *P*=0.06) (Fig. 1D) at 24 h after irradiation. Modulation of cell-surface CALR after irradiation was confirmed by immunofluorescence staining (Fig. 1E). FLASH RT increased the percentage of Calr^+^ TC1 cells at 24 h after irradiation compared with CONV RT or MOCK controls (Fig. 1F). Given that FLASH RT increased the expression of these two DAMPs, we also assessed expression of cGAS-STING pathway members *cGas (Mb21d1)*, Sting (*Tmem173)*, and *Ifna1* mRNA by using qRT-PCR. In TC1 murine cells, FLASH RT upregulated *cGas* mRNA (9 Gy, 2.0 vs. 0.9, *P*<0.001; 12 Gy, 1.9 vs. 1.0, *P*<0.001), *Sting* mRNA (9 Gy, 2.2 vs. 0.9, *P*<0.01; 12 Gy, 2.1 vs. 1.2, *P*<0.01), and *Ifna1* mRNA (9 Gy, 6.0 vs. 2.9, *P*<0.5; 12 Gy, 10.9 vs. 3.8, *P*<0.05) compared with CONV-RT (Fig. 1G-I). These findings indicate that FLASH RT may increase antitumor immune responses by increasing the expression of immunogenic DAMPs and cGAS-Sting.

**FIGURE 1.**
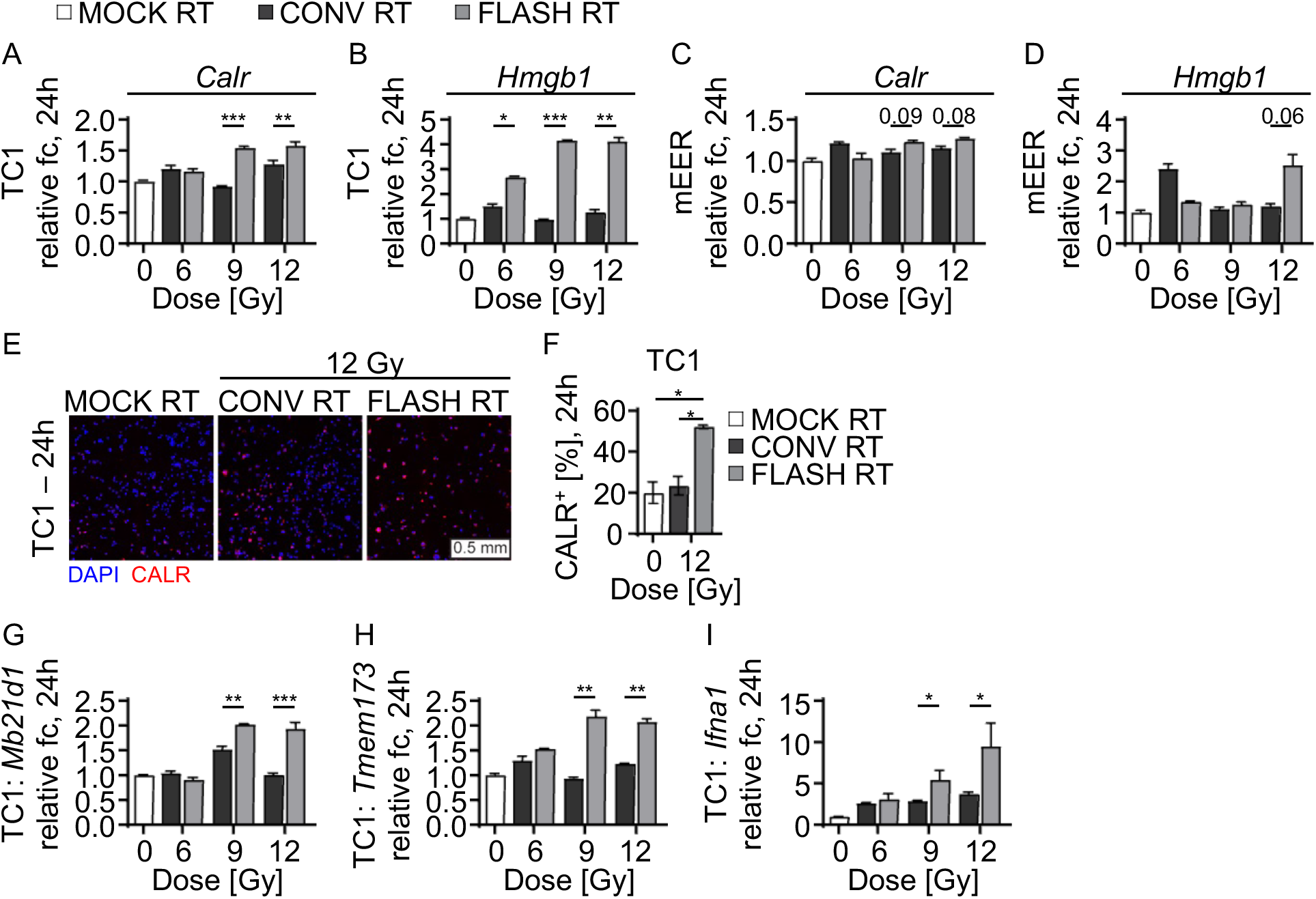
FLASH RT enhances DAMPs release and cGAS/STING upregulation in murine cancer cell lines. (A-D) Radiation-dose-dependent increases in mRNA expression of (A) CALR and (B) HMGB1 in TC1 cell lines at 24 h after irradiation and mRNA expression in (C) CALR and (D) HMGB1 in mEER cell lines at 24 h after irradiation. Gene expression is normalized to GAPDH and shown as relative fold-change (fc) from non-irradiated controls (2^-ΔΔ*Ct*^). (E) Representative images of CALR staining by immunofluorescence in TC1 cell lines at 24 h after irradiation. (F) Increased frequency of CALR^+^ TC1 cells from (E). DAPI (4’,6-diamidino-2-phenylindole) was used for counterstaining. Scale 0.5 mm. (G-I) Radiation-dose-dependent upregulation of (G) Mb21d1/cGas, (H) Tmem173/Sting, and (I) Ifna1 mRNA in TC1 cells at 24 h after irradiation. Representative data from three independent experiments. Mean fold change ± SEM. Paired, two-tailed Student’s *t* test \**P*<0.05, \*\**P*<0.01, \*\*\**P*<0.001. Abbreviations: cyclic GMP-AMP synthase (cGas), stimulator of interferon response cGAMP interactor 1 (Sting1).

### FLASH RT Spared Naïve and Activated Lymphocytes in Tumor-Draining Lymph Nodes

Mice bearing subcutaneous mEER tumors were irradiated daily with 9 Gy, given as FLASH or CONV RT, in 3 fractions to a total dose of 27 Gy; a tumor-targeting field was used to avoid directly irradiating the DLNs. Dosimetry output was monitored and recorded for each individual delivery for each mouse (Fig. 2A-D). At 5 days after the final RT fraction, we harvested peripheral blood samples and DLNs of tumor-bearing mice after treatment to 27 Gy by either FLASH RT or CONV RT. We assessed lymphocyte subsets in the tumor DLNs by using a 10-color spectral flow cytometric panel (Supplementary Table S2) and applied the gating strategy described in Supplementary Figure S1. Compared with CONV RT, FLASH RT led to increased numbers of CD8^+^ T cells (mean count 0.8×10^4^ vs. 1.7×10^4^; *P*<0.001) and CD4^+^ T cells (1.2×10^4^ vs. 2.3×10^4^; *P*<0.001); those increases were comparable to those in non-irradiated controls (Fig, 3). In the CD8^+^ T cell population, both naïve (CD62L^hi^ CD44^−^) and effector (CD62L^lo^ CD44^+^) T cells were increased (naïve, 5.5×10^3^ vs. 2.7×10^3^; *P*<0.001; effector, 2.2×10^3^ vs. 0.5×10^3^; *P*<0.001) in mice treated with FLASH RT relative to mice treated with CONV RT (Fig. 3B-D). In the CD4^+^ T-cell population, naïve (CD62L^hi^ CD44^−^) and effector (CD62L^lo^ CD44^+^) T cell subsets were also elevated (naïve, 0.8×10^3^ vs. 0.5×10^3^; *P*<0.05; effector, 1.1×10^4^ vs. 0.3×10^4^; *P*<0.01) in FLASH RT-treated mice (Fig. 3F-H). Although CD8^+^ and CD4^+^ T cells in the peripheral blood of FLASH RT-treated mice seemed to be higher than those in CONV RT-treated mice, those apparent differences did not reach statistical significance (Fig. 3I, J and Supplementary Figure S2). Therefore, these results indicate that FLASH RT spared lymphocytes in the tumor DLNs relative to CONV RT and approximated the lymphocyte levels of non-irradiated mice.

**FIGURE 2.**
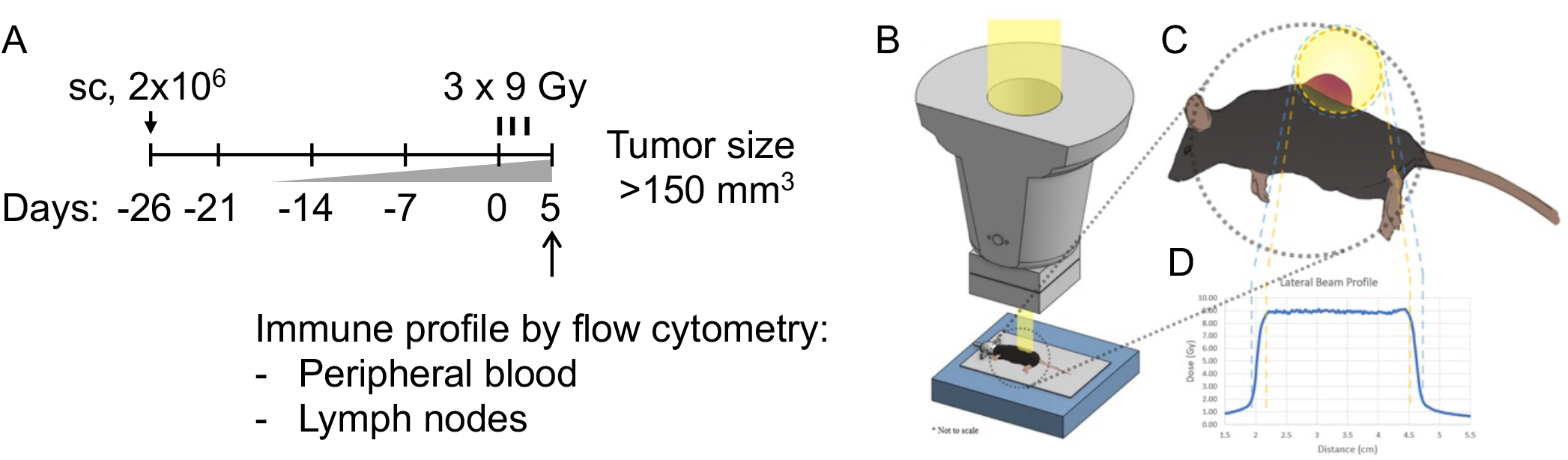
Delivery of hypo-fractionated FLASH- or CONV-RT in vivo. (A) Experimental design indicating timeline for tumor growth, radiation treatment (3 fractions of 9 Gy each), and endpoint analysis of immune cells from tumor DLNs and peripheral blood. (B) Illustration of the custom designed stereotactic jig setup indexed to the FLASH irradiator head. (C) Top view of mouse and subcutaneous tumor placement in relation to the radiation field and (D) lateral dose profiles.

**FIGURE 3.**
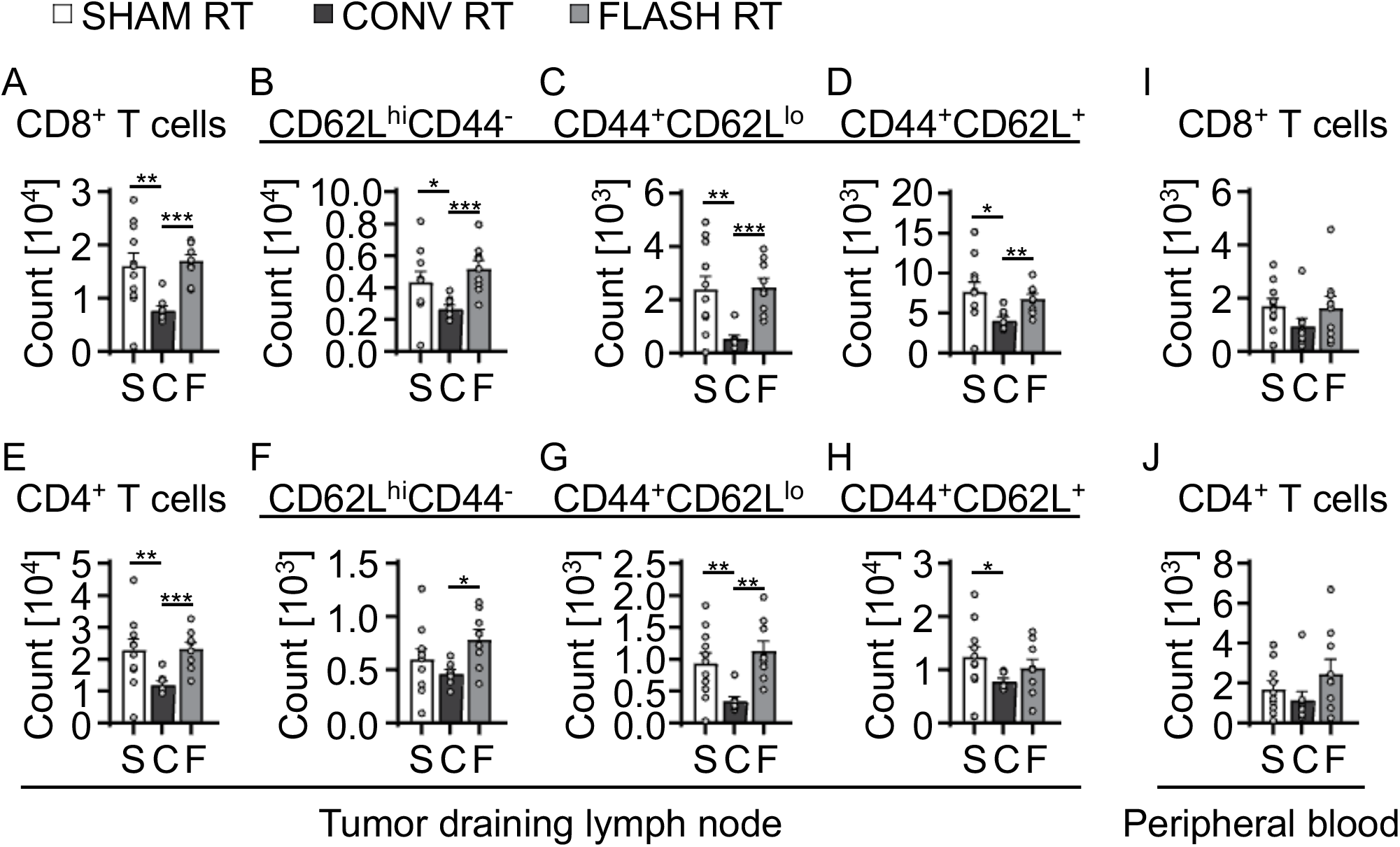
FLASH RT spares naïve immune T cells in tumor draining lymph nodes. Differences in (A) total CD8^+^ T cells and (B) CD4^+^ T cells isolated from tumor DLNs at 5 days after local subcutaneous tumor irradiation (27 Gy) quantified by flow cytometry. (C-E) Phenotypic characterization of (C) naive CD8^+^ cells (CD62L^hi^ CD44^+^), (D) effector CD8^+^ cells (CD44^+^ CD62L^lo^), and (E) central memory CD8^+^ cells (CD44^+^ CD62L^hi^) from (A). (F-H) Phenotypic characterization of (F) naive CD4^+^ T cells, (G) effector CD4^+^ T cells, and (H) central memory CD4^+^ T cells from (B). (I, J) Differences in circulating (I) total CD8^+^ T cells and (J) CD4^+^ T cells at 5 days after local subcutaneous tumor irradiation to 27 Gy. Gating strategies used for phenotypic characterization are shown in Supplementary Figure S1. Mean absolute counts ± SEM, biological repeats (n = 7-10). Paired, two-tailed Student’s *t* test \**P*<0.05, \*\**P*<0.01, \*\*\**P*<0.001.

### FLASH RT Increased Activated Macrophages in the Peripheral Blood

Because FLASH RT was associated with increased T-cell sparing, we assessed the effect of FLASH RT and CONV RT on myeloid cells in the peripheral blood. We observed that FLASH RT-treated mice had greater numbers of CD11b^+^ Ly6c^lo^ Ly6g^−^ macrophages (463.0 vs. 153.0, *P*=0.0561) and macrophages with upregulated MHC class II (195.0 vs. 61.0; *P*<0.05) than did CONV RT-treated mice, suggesting that FLASH spared activated macrophage populations (Fig. 4A, B). Similarly, greater counts of circulating CD11b^+^ Ly6c^high^ Ly6g^−^ monocytes (2.0×10^3^ vs. 1.1×10^3^; *P*<0.05) and MHC-II expressing monocytes (775.0 vs. 423.0; *P*<0.01) were detected in the peripheral blood after FLASH RT (Fig. 4C, D). In addition to the increased monocyte and macrophage counts, FLASH RT-treated mice exhibited significantly elevated numbers of circulating CD11c^+^ CD11b^−^ dendritic cells (801.4 vs. 360.6; *P*<0.05) and MHC-II upregulated dendritic cells (667.1 vs. 330.2; *P*<0.05) compared to CONV RT-treated mice, suggesting that FLASH RT better preserves dendritic cell populations in circulation (Fig. 4E, F). No differences were seen in dendritic cell populations in the tumor DLNs between FLASH RT- and CONV-RT-treated mice (Fig. 4E-H).

**FIGURE 4.**
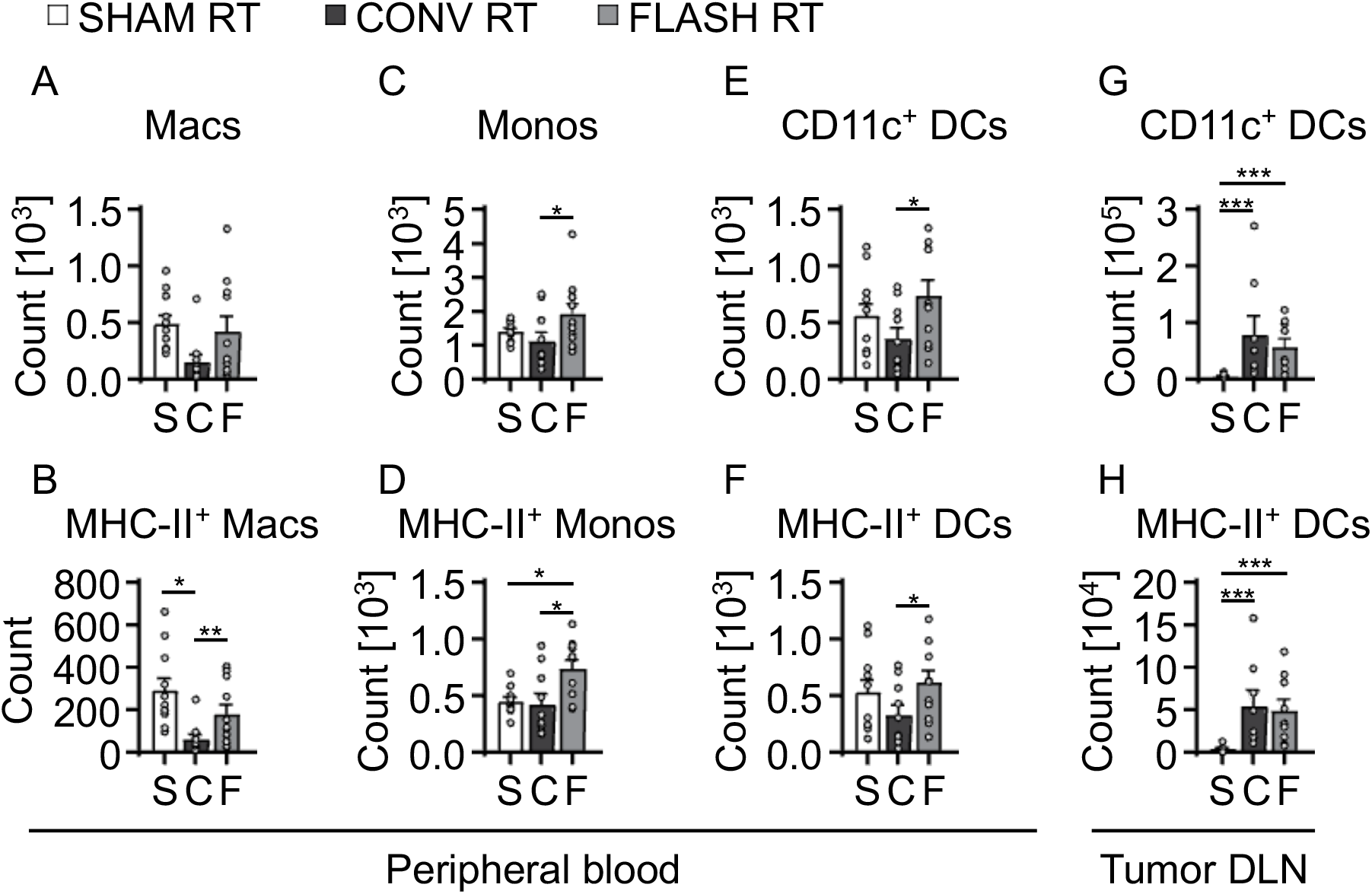
Increased activated macrophage and monocyte counts in peripheral blood after FLASH RT. (A) Total counts of macrophages (Macs) in peripheral blood from tumor-bearing mice at 5 days after tumor irradiation. (B) Counts of MHC-II upregulated Macs from (A). (C) Total counts of monocytes (Monos) in peripheral blood from tumor-bearing mice at 5 days after tumor irradiation. (D) Counts of MHC-II upregulated Monos from (C). (E) Total counts of dendritic cells (DCs) in peripheral blood from tumor-bearing mice at 5 days after tumor irradiation. (F) Counts of MHC-II+ activated DCs from E. (G) Total counts of DCs detected in tumor draining lymph nodes (DLNs) from tumor-bearing mice at 5 days after tumor irradiation. (H) Counts of MHC-II+ activated DCs from G. Gating strategies used for phenotypic characterization are shown in Supplementary Figure S1. Mean absolute counts ± SEM, biological repeats (n = 7-10). Paired, two-tailed Student’s *t* test \**P*<0.05, \*\**P*<0.01, \*\*\**P*<0.001.

### FLASH RT Spared Tumor-Infiltrating Immune Suppressive Cells

The immune checkpoint proteins programmed death-ligand 1 (PDL1) and PD1 are upregulated in irradiated tumors (5). To assess the effects of FLASH RT on the tumor immune microenvironment, we used multiplex immunofluorescence staining of FLASH RT- and CONV RT-treated tumors to assess immune checkpoint expression on tumor-infiltrating lymphocytes and macrophages (Fig. 5A, B). Compared with CONV RT-treated tumors, FLASH RT-treated tumors displayed 3.4-fold more CD8^+^ T cells (57.4 cells/mm^2^ vs. 16.7 cells/mm^2^, *P<*0.05; Fig. 5C). Furthermore, a greater percentage of cytotoxic T cells antigen-experience (PD1^+^ CD8^+^ T cells) was detected in tumors treated with FLASH RT compared with tumors treated with CONV RT (mean 31.0% vs. 6.2%, *P*<0.05; Fig. 5D). Although total numbers of F4/80^+^ macrophages were no different after FLASH vs CONV (Fig. 5E), FLASH RT-treated tumors had a greater percentage of tumor-associated macrophages (F4/80^+^PDL1^+^) compared with CONV RT-treated tumors (11.3% vs. 0.54%; *P*<0.05; Fig. 5F). Therefore, tumors treated with FLASH RT displayed increased tumor-infiltrating lymphocytes and increased expression of the checkpoint molecules PD1 and PDL1, suggesting antagonistic effects in the tumor microenvironment response.

**FIGURE 5.**
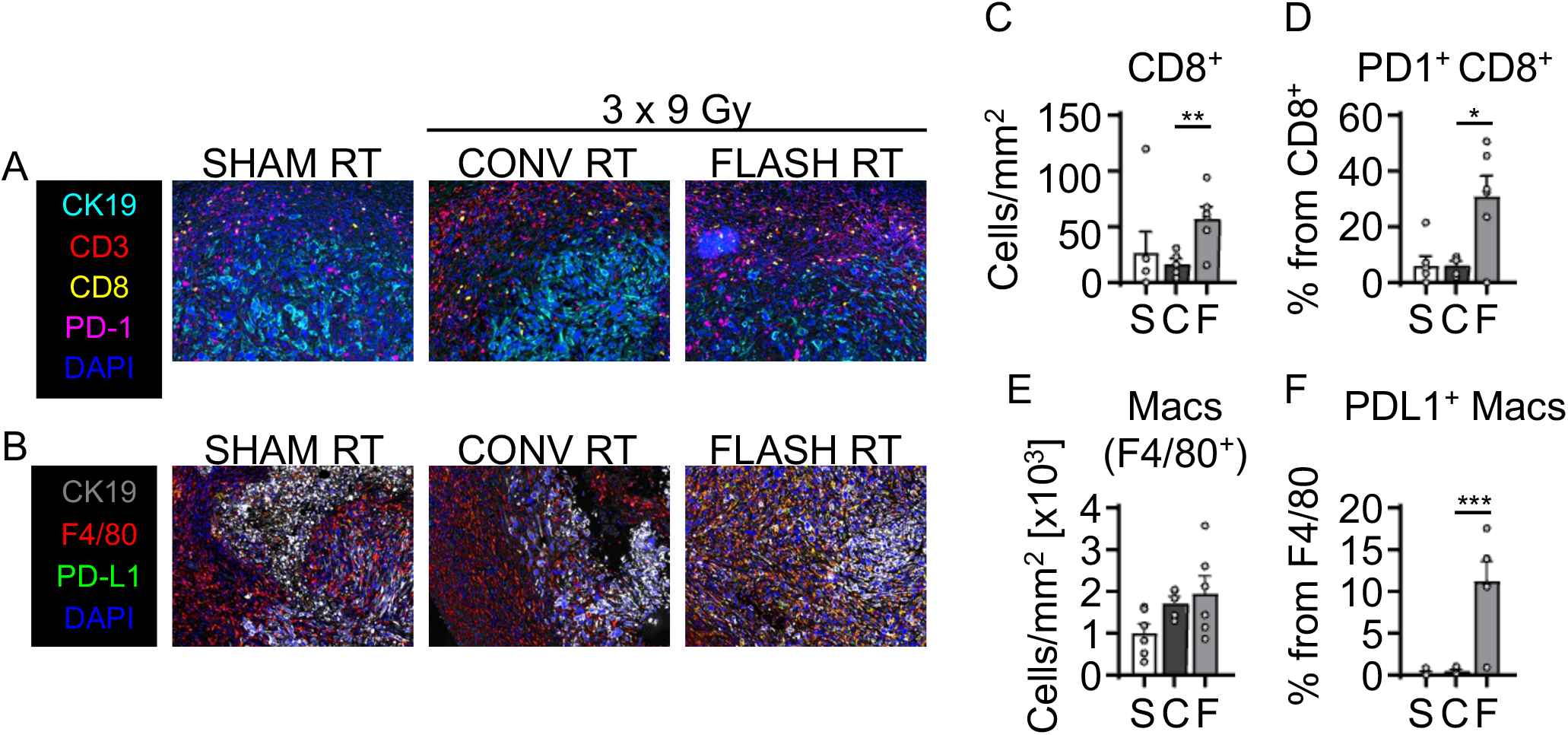
Increased tumor-infiltrating lymphocytes and tumor-associated macrophages with upregulated immune checkpoints in the tumor microenvironment of FLASH RT-treated tumors. (A, B) Representative regions of interest show (A) TILs and (B) tumor-associated macrophages by multiplex immunofluorescence staining at 5 days after irradiation. (C) Increased intratumoral CD8^+^ T cells observed in tumors treated with FLASH RT. (D) Higher frequency of TILs (PD1^+^ CD8^+^ T cells) in tumors at 5 days after FLASH RT. (E, F) Increased intratumoral (E) macrophages and (F) TAMs (PDL1^+^ Macs) at 5 days after FLASH RT. Gating strategies for phenotypic characterization a shown in Supplementary Figure S1. Representative of two independent experiments. Mean absolute counts, Mean percent ± SEM, biological repeats (n = 5-6 ROIs). Paired, two-tailed Student’s *t* test \**P*<0.05, \*\**P*<0.01, \*\*\**P*<0.001.

## Discussion

Here, we found that irradiation of syngeneic tumors with FLASH RT spared lymphocytes in tumors and in tumor-DLNs. To make these observations, we used a syngeneic tumor model with cells implanted in the mid-flank to minimize the overlap of the radiation field with the DLN. DLNs displayed greater numbers of naïve and activated CD8^+^ T cells when tumors were irradiated with FLASH RT compared with CONV RT. Furthermore, DLNs in mice irradiated with FLASH RT displayed greater numbers of naïve CD4^+^ T cells. This lymphocyte sparing by FLASH RT was also observed in tumor-infiltrating lymphocytes. Relative to CONV RT, FLASH RT also spared circulating macrophages and increased the numbers of activated macrophages, as indicated by increased MHC II expression. By contrast, dendritic cells were similarly increased in both FLASH RT- and CONV RT-treated mice. These findings are consistent with the increased expression of immunogenic proteins including DAMPs and cGAS-STING family members in cells irradiated with FLASH RT *in vitro*. Finally, FLASH RT was associated with increased immune checkpoint inhibitors in tumor-infiltrating lymphocytes and macrophages, suggesting that FLASH may also spare immunosuppressive mechanisms that limit antitumor immune responses. Thus, FLASH RT was associated with increased numbers of lymphocytes both in the tumor DLNs and in the irradiated tumors, suggesting that FLASH RT may better preserve antitumor immune responses than conventional RT strategies.

Previous groups have demonstrated that FLASH RT spares normal tissues including neural tissues, gastrointestinal epithelium, lung parenchyma, and others (12-18). By contrast, few if any reports have demonstrated immune cell sparing by FLASH RT. Part of the immune sparing, we observed may reflect a decrease in the fraction of blood that was within the radiation field. To this end, Galts et al. modeled the fraction of blood irradiated by proton beam scanning FLASH RT and CONV intensity-modulated proton therapy directed at brain tumors (30). Simulations predicted a nearly 5-fold decrease in peripheral blood irradiation with FLASH RT and reduced the depletion of circulating lymphocytes by 69%. Although reducing the total blood volume irradiated probably influenced lymphocyte counts in the current study, Venkatesulu et al. demonstrated that irradiation of heart tissues, where blood flow is high, UHDR did not reduce lymphopenia (31). However, Venkatesulu et al used lower mean dose rates (35 Gy/s) than those in the current study (270 Gy/s), so we cannot exclude that higher dose rates may have better lymphocyte sparing effects, especially in less well perfused tissues such as in peripheral tumors. The immune sparing we observed may also reflect reduced toxicity to lymphoid organs outside of the radiation field, because we directly observed increased numbers of lymphocytes in non-irradiated tumor DLNs. Similarly, Joseph et al. observed that total body integral dose correlated with worse lymphocyte counts after CONV RT (32). It is unlikely that the increase in lymphocytes in DLNs was solely due to increased sparing of lymphocytes from the peripheral blood because on average, CD8+ T cells spend 21 hours surveilling a single lymph node and thus probably remained within the lymph node compartment after RT (33). Therefore, our results suggest that lymphocyte sparing may occur both directly within the irradiated field and out of the field.

We also observed that FLASH RT increased the expression of immunogenic proteins including calreticulin, HMGB1 and cGAS-STING family members in irradiated cancer cells. By contrast, Shi et al. demonstrated that FLASH RT was less efficient in eliciting cGAS-STING activation in normal gastrointestinal epithelium (19). Katsuki et al. observed that carbon ion UHDR irradiation and CONV RT similarly increased calreticulin in melanoma cells (34). The induction of immunogenic proteins by FLASH RT may be cell line– and cell type–dependent. Regardless, FLASH RT may stimulate antitumor immune responses by increasing different immunogenic pathways.

Given our observation that FLASH RT was associated with increased PDL1, PD1, or both on immune cells, FLASH RT may uniquely benefit from the addition of immune checkpoint blockade. Focal, hypofractionated CONV RT, or stereotactic body RT, may lessen the immunosuppressive properties of RT in selected cases and synergize with immune checkpoint inhibitors (35,36). However, the existence and magnitude of this synergy were extremely variable across studies, and stereotactic body RT ultimately had lethal effects on resident immune cells and tumor-infiltrating lymphocytes. By contrast, FLASH RT both spared lymphocytes in the irradiated tumor and non-irradiated DLNs as well as upregulated immunosuppressive checkpoint molecules. Therefore, directly targeting the PD1/PDL1 axis may enhance the effectiveness of FLASH RT.

It is interesting to speculate that FLASH RT may also overcome the immune dysfunction associated with elective nodal irradiation (ENI). In both preclinical models and human cancers, ENI was associated with immune dysfunction in the tumor DLNs, as reflected by increased regulatory T cells and reduced effector T cell polarization (37,38). In preclinical models, ENI or surgical ablation of the tumor DLNs also reduced the efficacy of immunotherapy (37,39-42). These observations are consistent with the failure of immunotherapy to improve locoregional control or overall survival in cervical and head and neck cancers when immunotherapy is given concurrently with RT + ENI (8,43-46). By better sparing secondary lymphoid organs that are susceptible to RT, FLASH RT may rescue the immune dysfunction associated with ENI and better synergize with immunotherapies such as immune checkpoint inhibitors.

In summary, we observed that FLASH RT spared lymphocytes in tumor DLNs and irradiated tumors, including both naïve and effector T cell populations. In tumors, FLASH RT was also associated with an increase in the immune checkpoint molecules PD1 on CD8^+^ T cells and PDL1 on macrophages, indicating that FLASH RT may also spare immunosuppressive mechanisms that inhibit radiation-induced antitumor immunity. Consequently, FLASH RT may uniquely benefit from the addition of immune checkpoint inhibitors, given the lymphocyte-sparing effects that may be blunted by the preservation of immunosuppressive mechanisms within the tumors. Overall, these results indicate that FLASH RT enhanced antitumor immune responses, an effect that could be augmented with immunotherapies.

## Authors’ Contributions

**F.R. Saenz:** Conceptualization, data curation, formal analysis, investigation, methodology, project administration, validation, visualization, writing, reviewing & editing. **B. Velasquez:** Data curation, investigation, methodology, validation, writing. **T. Waldrop:** Resources. **E. Aguilar:** Data curation, investigation, resources. **K.R. Cox:** Resources. A. Delahoussaye: Resources. **C. Laberiano-Fernandez:** Data curation, formal analysis, methodology. **L. Campos Clemente:** Data curation, formal analysis, methodology. **L. Connell:** Resources. **N. Mims:** Resources. **D. Neill:** Resources. **E.R. Parra:** Methodology, project administration. **K. Clise-Dwyer:** Data curation, formal analysis, methodology. **E. Schüler:** Conceptualization, formal analysis, methodology, funding acquisition, project administration, supervision, writing, reviewing & editing. **M.T. Spiotto:** Conceptualization, formal analysis, methodology, funding acquisition, project administration, supervision, writing, reviewing & editing.

## Acknowledgements

We thank Christine F. Wogan, MS, ELS, of MD Anderson’s Division of Radiation Oncology, for editorial contributions to several drafts of this article. Research reported in this publication was supported in part by the National Cancer Institute of the National Institutes of Health under Award Number R01CA266673, by the University Cancer Foundation via the Institutional Research Grant program at MD Anderson Cancer Center, by a grant from MD Anderson’s Division of Radiation Oncology, by Cancer Center Support Grant P30 CA016672 (Research Animal Support Facility and the Advanced Cytometry and Sorting Core Facility) from the National Cancer Institute of the National Institutes of Health, to The University of Texas MD Anderson Cancer Center, and by training fellowships from UTHealth Houston Center for Clinical and Translational Sciences Program (Grants No. TL1 TR003169; T32 TR004905). The content is solely the responsibility of the authors and does not necessarily represent the official views of the National Institutes of Health nor of the Cancer Prevention and Research Institute of Texas.

## Notes

### Competing Interest Statement

The authors have declared no competing interest.

