## Supplementary materials for "FLASH radiotherapy spares lymphocytes in tumor-draining lymph nodes and increases infiltration of immune cells in tumors"

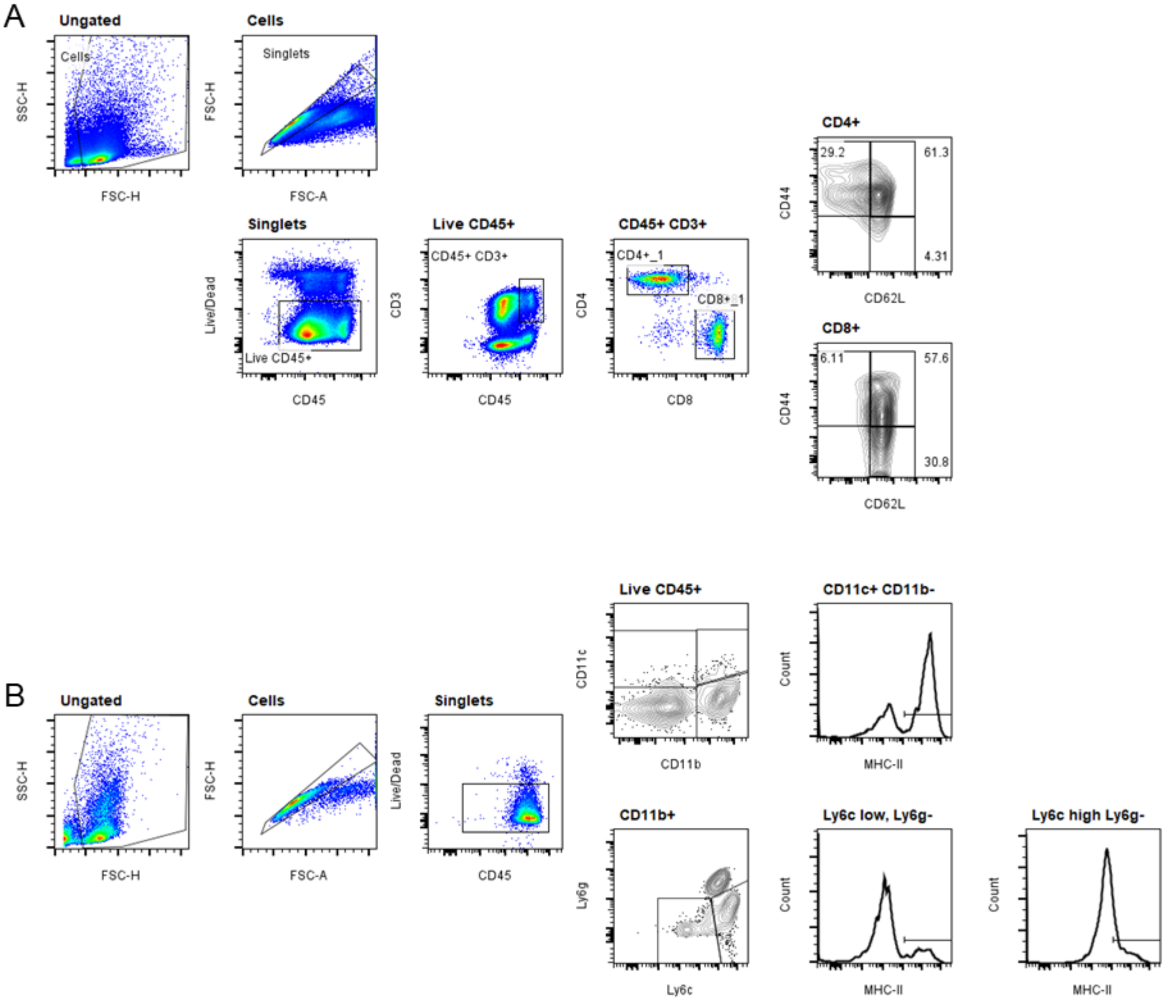


**Supplementary Figure S1. Gating strategy for the characterization of immune cells by spectral flow cytometry.**

(A) Lymphoid cells gating strategy: CD45$^+$ cells were gated on CD3$^+$ cells to identify CD4$^+$ and CD8$^+$ T cells. Effector, memory, and naive T cells were further characterized based on CD44 and CD62L expression. (B) Myeloid cells gating strategy: Antigen-presenting cells were identified by gating on MHC Class II and CD45 expression. Dendritic cells were identified based on CD11c expression. CD11b$^+$ cells were further characterized by Ly6g and Ly6c expression to identify granulocytes (Ly6g$^+$ Ly6c$^+$), macrophages (Ly6g$^-$ Ly6c$^{lo}$), and monocytes (Ly6g$^-$ Ly6c$^{hi}$).

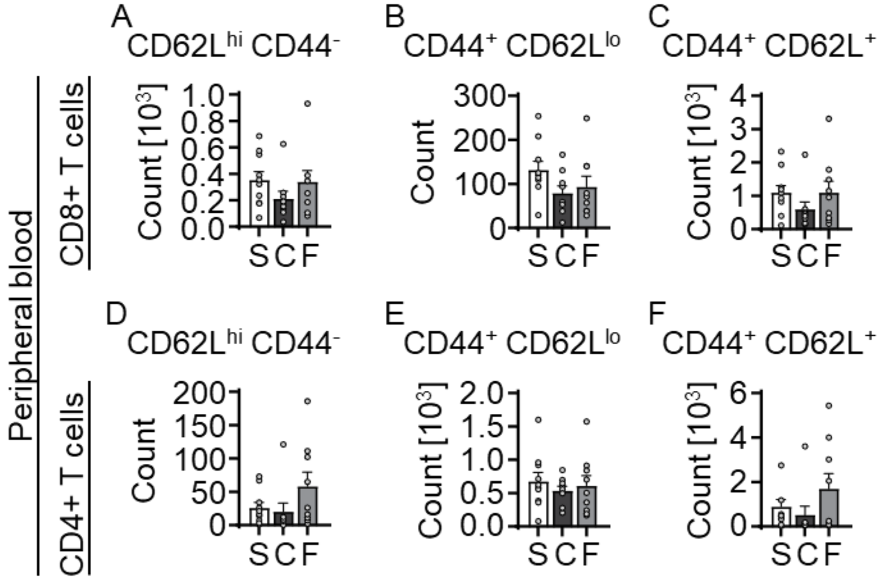


**Supplementary Figure S2. Phenotypic characterization of circulating lymphocytes after FLASH RT.**

Phenotypic analysis of CD8+ and CD4+ T cell subsets in peripheral blood 5 days after irradiation. (A–C) Flow cytometry analysis of CD8+ T cells, showing (A) Naive (CD62L$^{hi}$ CD44$^{-}$), (B) Effector (CD44$^{+}$ CD62L$^{lo}$), and (C) Central memory (CD44$^{+}$ CD62L$^{+}$) populations. (D–F) Quantification of CD4+ T cell subsets, including (D) Naive (CD62L$^{hi}$ CD44$^{-}$), (E) Effector (CD44$^{+}$ CD62L$^{lo}$), and (F) Central memory (CD44$^{+}$ CD62L$^{+}$) populations. Phenotypic characterization was performed using the gating strategy detailed in Supplementary Figure S1. Mean absolute counts ± SEM, biological repeats (n = 7-10).

**Supplementary Table S1.** Physical beam parameters measured using Gafchromic film and beam current transformers during cell irradiations.

| | Total dose (Gy) | Number of pulses (N) | Dose per pulse (Gy) | Mean Dose Rate (Gy/s) | Irradiation time (s) | Intrapulse dose rate (MGy/s) | Pulse width (µs) | Pulse repetition frequency (Hz) |
|---|---|---|---|---|---|---|---|---|
| FLASH | 6.3 ± 0.07 | 2 | 3.2 ± 0.04 | 284 ± 4 | 0.02 | 2.6 | 1.2 | 90 |
| CONV | 6.1 ± 0.04 | 437 | 0.01 | 0.4 | 15 | 0.01 | 1.2 | 30 |
| FLASH | 9.2 ± 0.10 | 3 | 3.1 ± 0.04 | 276 ± 4 | 0.03 | 2.6 | 1.2 | 90 |
| CONV | 9.1 ± 0.06 | 655 | 0.01 | 0.4 | 22 | 0.01 | 1.2 | 30 |
| FLASH | 12.5 ± 0.14 | 4 | 3.1 ± 0.04 | 281 ± 4 | 0.04 | 2.6 | 1.2 | 90 |
| CONV | 12.1 ± 0.08 | 873 | 0.01 | 0.4 | 29 | 0.01 | 1.2 | 30 |

**Supplementary Table S2.** Physical beam parameters measured using Gafchromic film and beam current transformers during animal irradiations.

| | Total dose (Gy) | Number of fractions | Dose per fraction (Gy) | Number of pulses (N) | Dose per pulse (Gy) | Mean Dose Rate (Gy/s) | Irradiation time (s) | Intrapulse dose rate (MGy/s) | Pulse width (µs) | Pulse repetition frequency (Hz) |
|---|---|---|---|---|---|---|---|---|---|---|
| FLASH | 27.5 ± 0.31 | 3 | 9.2 ± 0.12 | 3 | 3.1 ± 0.04 | 275 ± 4 | 0.03 | 0.97 | 3.2 | 90 |
| CONV | 27.1 ± 0.19 | 3 | 9.1 ± 0.14 | 1317 | 0.01 | 0.2 | 44 | 0.01 | 1.2 | 30 |

**Supplementary Table S3.** Comprehensive list of immune panels, flow cytometry antibodies, and working concentrations for staining of 0.2–3.0 × $10^6$ cells in 0.1 mL buffer.

| Marker | Fluorophore | Company | Catalog # | Clone | WC [0.1 mL] (0.2 - 3x$10^6$ cells) | Panel |
|---|---|---|---|---|---|---|
| Live/Dead | Ghost UV450 | Cytek Biosciences | SKU 13-0868-T500 | NA | 1/1000 | Lymph, Myeloid |
| CD4 | BUV737 | BD Biosciences | 612844 | RM4-5 | 0.2 ug/ml | Lymph |
| CD69 | BV421 | BioLegend | 104528 | H1.2F3 | 0.5 ug/ml | Lymph |
| CD44 | BV650 | BioLegend | 103049 | IM7 | 0.5 ug/ml | Lymph |
| PD-1 | BV786 | BioLegend | 135225 | 29F.1A12 | 1.0 ug/ml | Lymph |
| CD62L | PerCP-Cy5.5 | BioLegend | 104432 | MEL-14 | 1.0 ug/ml | Lymph |
| PD-L1 | PE | BioLegend | 124308 | 10F.9G2 | 0.5 ug/ml | Lymph |
| CD8 | PE-Cy7 | BioLegend | 100722 | 53-6.7 | 0.2 ug/ml | Lymph |
| CD3 | APC | BioLegend | 100236 | 17A2 | 0.4 ug/ml | Lymph |
| CD45 | APC-Cy7 | BioLegend | 103116 | 30-F11 | 0.2 ug/ml | Lymph, Myeloid |
| Ly6c | BUV737 | BD Biosciences | 755201 | HK1.4.rMAb | 0.2 ug/ml | Myeloid |
| CD11c | BV421 | BioLegend | 117330 | N418 | 0.25 ug/ml | Myeloid |
| CD172a | BV510 | BioLegend | 144032 | P84 | 2.0 ug/ml | Myeloid |
| CD86 | BV605 | BioLegend | 105125 | PO3 | 0.5 ug/ml | Myeloid |
| I-A/I-E | BV785 | BioLegend | 107645 | M5/114.15.2 | 0.2 ug/ml | Myeloid |
| Ly6g | PerCP-Cy5.5 | BioLegend | 127616 | 1A8 | 0.4 ug/ml | Myeloid |
| XCR1 | PE | BioLegend | 148204 | ZET | 0.7 ug/ml | Myeloid |
| CD163 | PE-dazzle594 | BioLegend | 156710 | S15049F | 0.5 ug/ml | Myeloid |
| CD11b | PE-Cy7 | BioLegend | 101216 | M1/70 | 0.2 ug/ml | Myeloid |
| F4/80 | APC | BioLegend | 123116 | BM8 | 0.5 ug/ml | Myeloid |
| CD16/32 | Blocking | BioLegend | 101302 | 93 | 1.0 ug/ml | Lymph, Myeloid |

**Supplementary Table S4.** Multiplex immunofluorescence mouse panel 1B, list of antibodies, and working concentrations.

| Target molecule | Description | Clone | Vendor | Cat# | Dilution Factor |
|---|---|---|---|---|---|
| CK19 | Tumor cells | TROMA-III-S | DSHB | AB2133570 | 1:20 |
| CD8 alpha | CD8+ T cells | D4W2Z | CELL SIGNALING | 98941S | 1:200 |
| F4/80 | Macrophages | D2S9R | CELL SIGNALING | 70076S | 1:200 |
| CD4 | CD4+ T cells | D7D2Z | CELL SIGNALING | 25229S | 1:100 |
| CD3 epsilon | T lymphocyte | D4V8L | CELL SIGNALING | 99940S | 1:100 |
| PD-L1 | APCs, B cells, Tumor cells | D5V3B | CELL SIGNALING | 64988S | 1:100 |
| PD1 | Activated T cells, B cells, monocytes | D7D5W | CELL SIGNALING | 84651S | 1:50 |
| CD19 | B cells | D4V4B | CELL SIGNALING | 90176S | 1:100 |
| DAPI | Nucleus | | AKOYA BIOSCIENCE | FP1490 | 4drops 10X/ml |
